# Multiple Imputation Approaches Applied to the Missing Value Problem in Bottom-up Proteomics

**DOI:** 10.1101/2020.06.29.178335

**Authors:** Miranda L. Gardner, Michael A. Freitas

## Abstract

Analysis of differential abundance in proteomics data sets requires careful application of missing value imputation. Missing abundance values vary widely when performing comparisons across different sample treatments. For example, one would expect a consistent rate of “missing at random” (MAR) across batches of samples and varying rates of “missing not at random” (MNAR) depending on inherent difference in sample treatments within the study. The missing value imputation strategy must thus be selected that best accounts for both MAR and MNAR simultaneously. Several important issues must be considered when deciding the appropriate missing value imputation strategy: (1) when it is appropriate to impute data, (2) how to choose a method that reflects the combinatorial manner of MAR and MNAR that occurs in an experiment. This paper provides an evaluation of missing value imputation strategies used in proteomics and presents a case for the use of hybrid left-censored missing value imputation approaches that can handle the MNAR problem common to proteomics data.

## INTRODUCTION

Improvement in mass spectrometry instrumentation and methodologies along with a growing interest in the exploration of multi-omics, incorporating proteomics with genomics for targeted therapeutics, have led to generation of large, high-density datasets and introduced a new confounding factor in data analysis: missing values [1–4]. Missing values in mass spectrometry-based proteomics data analysis can range from 5 – 50% in any given replicate for observed peptides abundances. Two approaches to deal with missing values are 1) removing peptides/proteins that have insufficient samples for analysis or 2) imputing values as placeholders for the missing values [5]. The former approach may be acceptable when analyzing a small number of samples with similar proteomic profiles and few missing values. However, the latter imputation approach, when appropriately applied, can avoid unnecessarily excluding data from an analysis.

The most common sources of missing values in proteomics experiments are 1) the biology and/or technical sample preparation, 2) actual presence below the instrument’s limit of detection (LOD) threshold and 3) presence above the LOD but error in data preprocessing [6]. Furthermore, missing values in proteomics data can be classified into one of three categories: missing completely at random (MCAR), missing at random (MAR) or missing not at random (MNAR) [7]. MCAR is independent of the data and observed values, much as it’s name suggests, and is likely to occur across the entire distribution of data. This type of missingness originates from inaccurate instrumentation or oversight in experimental sample preparation. MAR covers a wider range than MCAR and the missingness is due to conditional dependency on observed values. This class of missing values can arise when the peptide sequence is mapped incorrectly or software erroneously assigns shared peptides to precursors leading to misidentification in some samples and missing values in others.

MAR methods considered for this study include *k* nearest neighbors (*k*NN), singular value decomposition (SVD) and maximum likelihood estimate (MLE). The *k*NN algorithm imputes missing values to the gene of interest from genes with similar expression profiles. The missing value is estimated from a weighted average of the *k* closest genes to the gene of interest [8]. The SVD method converts the dataset to eigengenes, a principle component gene expression matrix. This algorithm fills in missing values with row averages and performs consecutive iterations with eigengene regression until the total change in the matrix falls below 0.01 [8]. MLE assumes that data is a function of an unknown parameter θ. The imputed value is a random draw of the MLE of θ that maximizes the probability of the observed data [9–11].

The MAR examples of missingness contrast a non-ignorable case of missing values where the missing values arise as a direct relationship with the data, MNAR. MNAR may result from experimental effects in proteomics such as 1) enzyme miscleavages, 2) true presence/absence (as seen in immunoprecipitation when comparing a treatment to an immunoglobulin control), and 3) instrumentation effects (dynamic range or LOD occurring when peptide measurements are low in abundance compared to background noise or constitute low ionization efficiency). Because the missingness is influenced by the low abundant nature of these values, this category of missing values is considered left-censored where the distribution of values (if present in the data) would fall on the left tail of the total observations in the data set.

Non-ignorable MNAR methods considered for this study include deterministic minimum (MinDet), probabilistic minimum (MinProb) and quantile regression imputation of left-censored data (QRILC). MinDet replaces each missing value with the smallest detectable intensity across the entire data set or observed within each sample [12, 13]. Similar to MinDet, MinProb also replaces missing values, but data only after the data is first centered on the MinDet value and the replacement is a random draw from the Gaussian distribution [11, 14]. The QRILC approach utilizes quantile regression to construct a truncated distribution from the left-most tail of the data and the missing values are then replaced with random draws from this reduced allotment [15].

Current literature on missing values imputation has explored the type of imputation (single / multiple), nature of missingness (MAR, MCAR or MNAR applied across entire datasets), statistical algorithms to determine differentially expressed proteins (DEP) (reviewed further in [5, 16–19]) and development of software tools to visualize the results (discussed further in [20–24]). Single imputation strategies work well with datasets that are very similar in nature (low number of missing values), simulated missing values and very large time course studies containing multiple time points and biological replications [11, 25]. However, as the number of missing values increases, it has been suggested that a single value estimate is not capable of generating the missing value accurately or capturing the variability. Additionally, a value too small or too large from the “true value” will heavily influence the downstream statistical analysis [26, 27].

The multiple imputation (MI) approach addresses the single value estimate concern by performing consecutive iterations of a chosen method to generate *m* imputed datasets, followed by a selection step that combines every *m* imputations into a final dataset for downstream analysis. These methods work well with small datasets but the imputation method and modeling approach are data dependent, sensitive to parameter selection and may need further optimization for effective implementation [26, 28, 29]. In contrast, multiple imputation in multi-factor analysis (MI-MFA) has been successfully applied to large datasets without *a priori* knowledge of the missingness. However, the MI-MFA approach assumes that missing values are MAR, requires good donor-recipient matches and may introduce bias if the sample size is too small yielding too few donors in the donor pool [27].

As it is often difficult to determine the main contributor to missing values, approaches that combine MAR and MNAR methods are important to consider. Here, we provide a comparison of several different combinations of workflow imputation methods available via the imputeLCMD package in R [11, 30] and offer insight into the most appropriate workflow and method for handling the missing data problem in proteomics. We further demonstrate that across increasing number and type of missingness, the MNAR/MAR MI SFI-hybrid approach consistently outperformed all other methods, as evidenced by the consistent range of logFC protein expression values and *q*-value significance in DEP comparisons.

## EXPERIMENTAL PROCEDURES

### Proteomic Datasets

#### Glucose Deprivation

This dataset originated from Lee *et al* [31] in which triple negative breast cancer cell line MDA-MB-468 was exposed to high glucose or glucose deprivation for 48 hours. Samples were prepped for bottom-up proteomics and mass spectra collected on a Q Exactive™ (Thermo Fisher Scientific) operated in data-dependent acquisition (DDA) mode. The dataset consisted of 2,525 proteins and the authors identified 681 DEPs (*p*-value < 0.01). The RAW Thermo files were downloaded from the PRoteomics IDEntifications database (PRIDE): PXD013966.

#### Pluripotent Cell Differentiation

The NTERA2 (NT2) dataset contained proteomics obtained after IP of Polycomb complex subunits EZH2, SUZ12 and IgG control [32]. IP of nuclear lysates were performed on undifferentiated and NT2 cells and NT2 cells treated for 8 days with retinoic acid to induce differentiation. Samples were prepped for bottom-up proteomics and mass spectra collected on a Q Exactive™ (Thermo Fisher Scientific) operated in DDA mode. After subtracting IgG background, 366 candidate EZH2-interactors and 191 candidate SUZ12-interactors were identified (FDR threshold = 0.05). The RAW Thermo files were downloaded from the PRIDE database: PXD004462.

### Database searching and label-free quantitation (LFQ)

Mass spectra from both sample sets were searched with the OpenMS platform and X!Tandem search engine against a reviewed UniProt human proteome (05/19/2019) containing the cRAP and MaxQuant contaminant FASTAs with the following parameters: full trypsin digest, 2 missed cleavages, variable modifications (oxidation of methionine +15.99491, carbamidomethyl of cysteine +57.02146), precursor (MS1) mass tolerance 20 ppm and fragment (MS2) mass tolerance 0.02 Da. PSM rescoring was completed with Percolator and protein inference was performed with FIDO across all samples, setting peptide and protein FDR to 0.05.

### Data processing, imputation, differential expression analysis and top protein lists

Prior to imputation and differential expression analysis, the MAR functions (*k*NN, MLE and SVD) were altered slightly to allow the random seed generator to freely sample imputation values within consecutive iterations of that method. Additionally, the first element of the model selector that flags the data as a ‘1’ for MAR or ‘0’ for MNAR was replaced with a vector of ‘1s’ so that all MAR values would be imputed. Data was processed via one of three workflows: Filter-Imputation-Selection (FIS or Entire Dataset Imputation Method), Filter-Selection-Imputation (FSI or Treatment Imputation Method) or Selection-Filter-Imputation (SFI Method) (**Fig. 1**). During selection, data columns were first grouped by sample or treatment type and then chosen for the appropriate pair-wise comparisons. Lowly expressed proteins were removed from the data set if the minimum number of observations was < 3 and total intensity was < 2e15. Missing values were imputed as all MAR (*k*NN, MLE or SVD), all MNAR (MinDet, MinProb or QRILC) or MAR/MNAR (*k*NN and QRILC) for 25 consecutive iterations. For the MAR methods, the model selector was converted to disallow MNAR imputation values from occurring. The typical model selector was utilized during the MAR/MNAR strategy but the imputation was performed separately for each treatment or group of data. Therefore, we have introduced this as a hybrid method. Quantile normalization was performed and limma was used to determine DEPs. Datasets from each method were then rank-ordered by *q-value* mean to generate the top 200 protein lists (**Fig. 3 and Supp. Fig. 5 and 9**). Spread plots were generated from the same rank-ordered lists: the data was filtered for the top 10 proteins from each method and merged into one common list, retaining all unique identifications and removing duplications (**Figs. 4, 5, Supp. Fig. 6, and Supp. Tables 2-4**). Known proteins in the canonical PRC2 complex (AEBP2, EED, EZH2, JARID2, PCL (MTF2 in this case), RbAp46 and SUZ12) were scored according to significance position across all methods. After totaling the methods in both IPs, final ranking was SFI-hybrid, MinDet, MinProb, QRILC, SVD, *k*NN and MLE (**Fig. 5D**). All source code for this investigation was completed with various packages using R (version 3.6.2) [33] and is available in supplemental material (**Supplemental File 1**).

**Figure 1.**
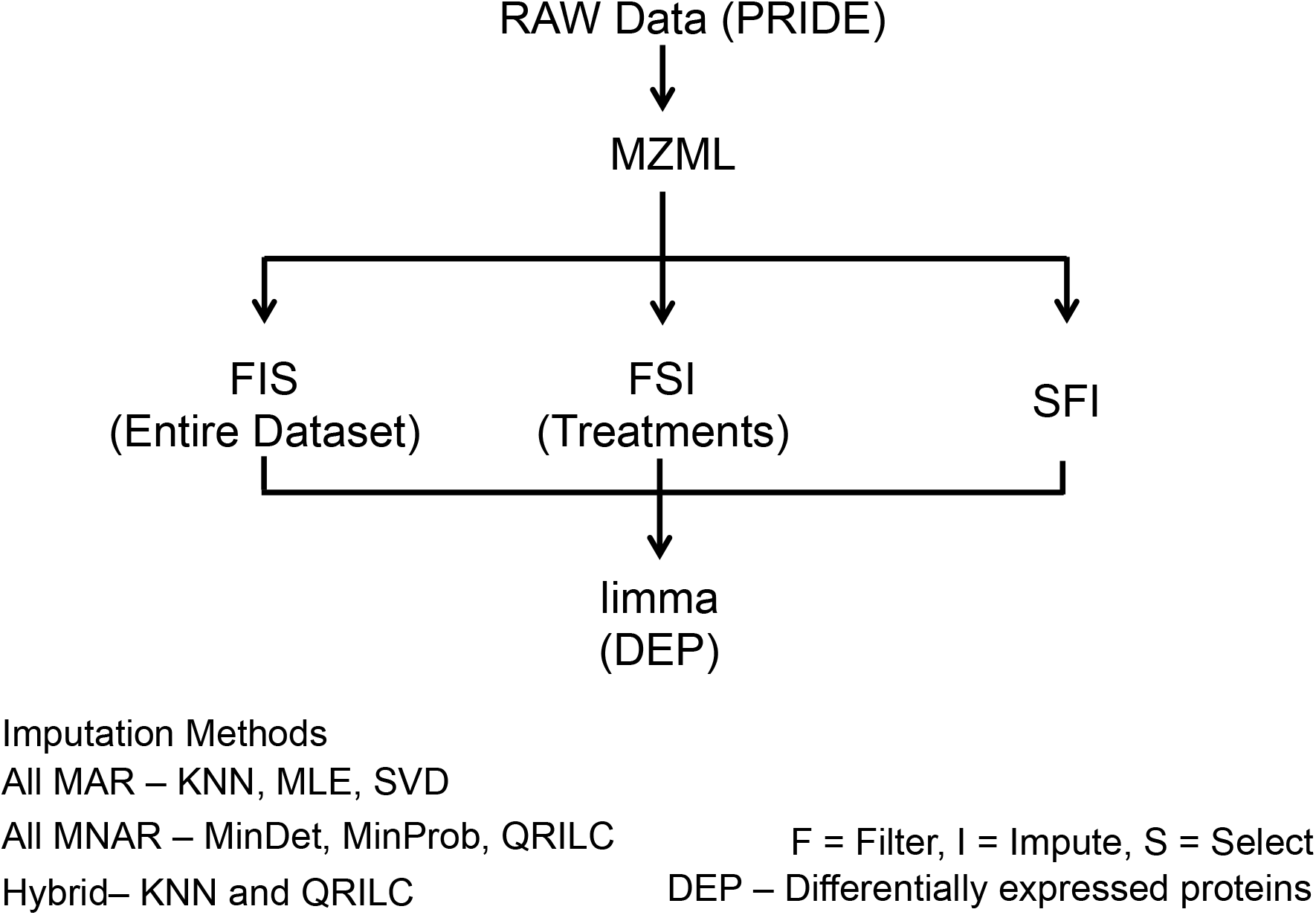
Data processing workflow. RAW proteomics data files were downloaded from PRIDE, converted to MZML prior to searching in OpenMS with X!Tandem search engine, processed via one of three methods (FIS, FSI or SFI) consisting of one of the 7 imputation methods (performed 25 consecutive iterations) and quantile normalized before utilizing limma to determine DEPs.

## RESULTS

### Imputation with small dataset of similar proteomic profiles

The PXD013966 dataset consisted of triple negative breast cancer cells MDA-MB-468 exposed to high glucose (n = 3) or glucose deprivation (n = 3) for 48 hours prior to sample preparation and analysis for bottom-up mass spectrometry. This whole cell proteomics experiment is representative of a small dataset with treatment and control in biological triplicate where most of the proteins are present and changes in protein expression levels are detected. The number of missing peptide values ranged from 8 – 15% across samples with a similar protein expression as evidenced by the distribution pattern of observed and missing peptides (**Fig. 2A**).

**Figure 2.**
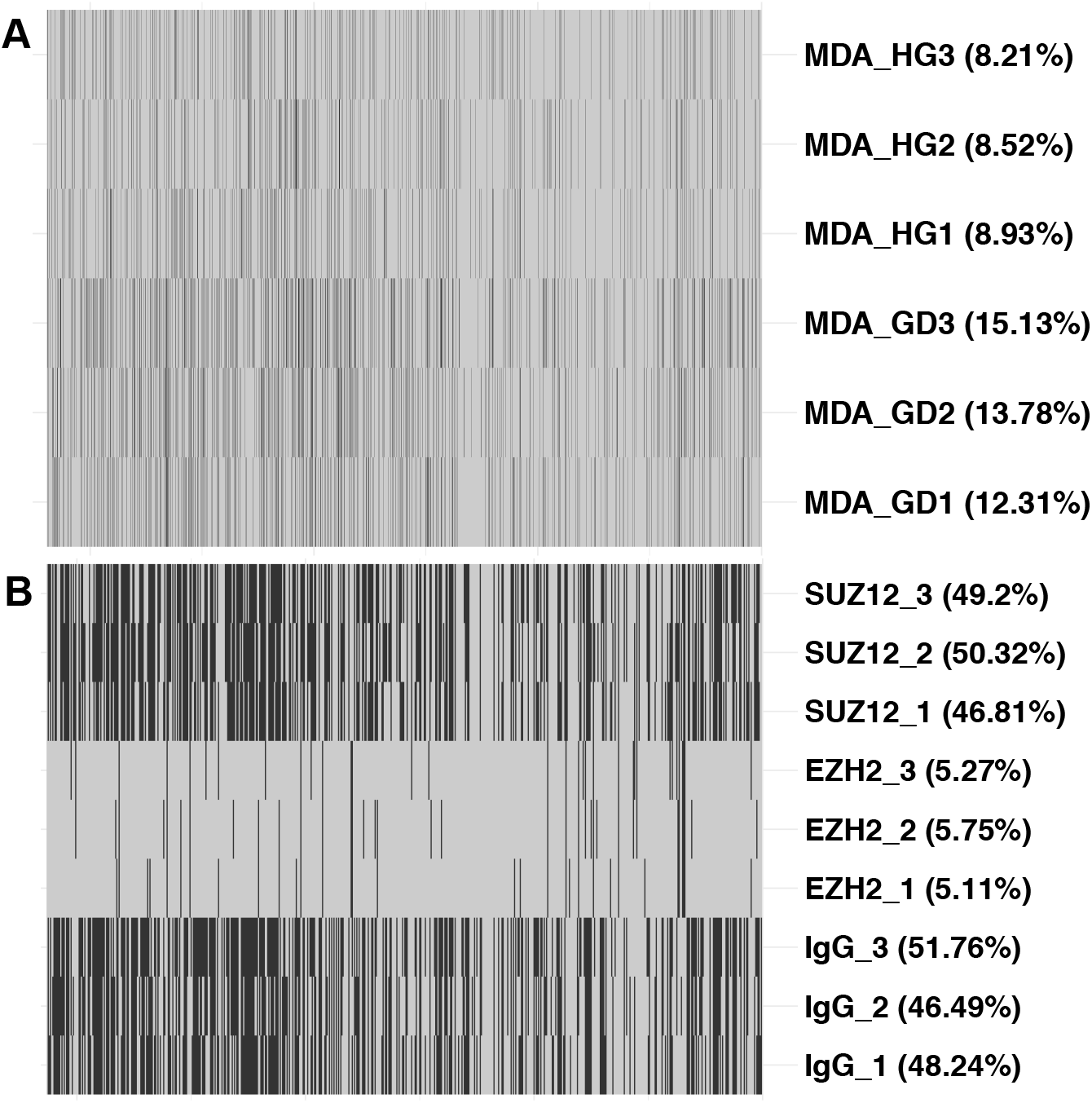
Distribution of number of missing values in the two datasets analyzed for this manuscript. The MDA data (**A**) is very similar in terms of the number of missing values (depicted as black lines) relative to observed values (depicted as gray lines) while the IP data (**B**) is more representative of a typical bottom-up proteomics experiment.

Following removal of contaminants, reverse decoys and lowly expressed proteins, we identified 3,165 total proteins. Missing values did not influence the range of logFC protein expression values with any of the methods **(Supp. Fig. 1)**. After performing 25 consecutive imputations, we ranked each imputation method DEP list by *q*-value mean. Using a threshold of 0.05 for significance cut-off, we observed the following DEP lists: 1,186 significant proteins when using *k*NN, 1,180 for MLE, 1,106 for SVD, 1,299 for MinDet, 1,312 for MinProb, 1,292 for QRILC and 1,669 for SFI-hybrid.

We further classified the top 200 DEPs from each method into the number and type of missingness: missing 0, 1, 2, or 3 values in one sample group (0, 1, 2, 3) or missing any combination of 2, 3 or 4 values in both groups (B2, B3, B4). With the exception of the SFI-hybrid method, the majority of the proteins determined to be significant were those with all observations present (**Fig. 3**). We discovered the SFI-hybrid approach demonstrated the most reproducibility in terms of standard deviation of logFC expression and maintaining *q*-value significance (**Fig. 4, Supp. Figs. 1 and 2**).

**Figure 3.**
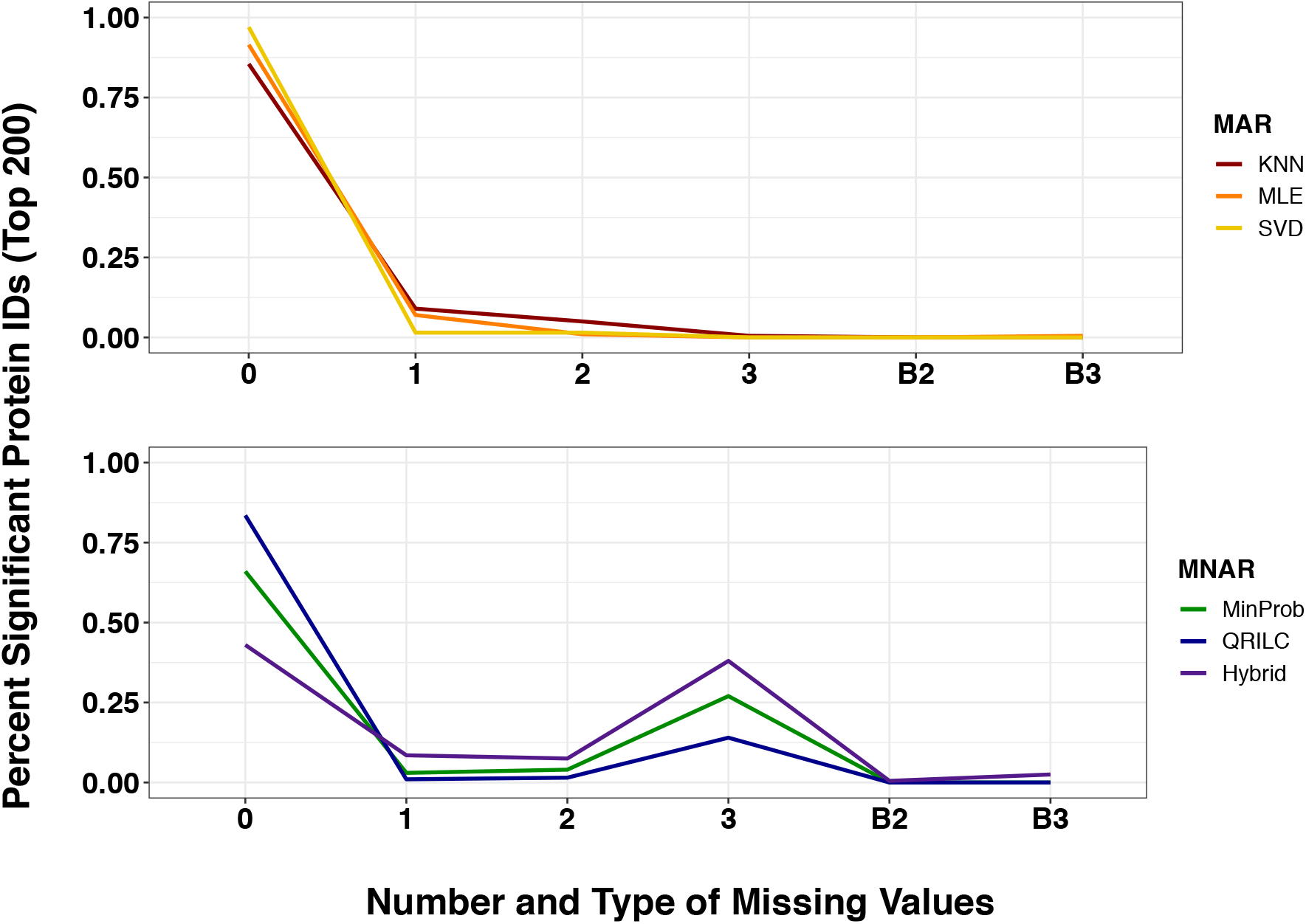
Frequency of missingness across MAR **(Top Panel)** and MNAR **(Bottom Panel)** imputation methods with the top 200 significant proteins in MDA data. After 25 consecutive iterations, each imputation dataset was rank-ordered by *q*-value mean. The top 200 proteins from each method was binned according to the type of missingness: 0, 1, 2 or 3 missing values in one sample group or combination of 2 or 3 in both groups (B2, B3).

**Figure 4.**
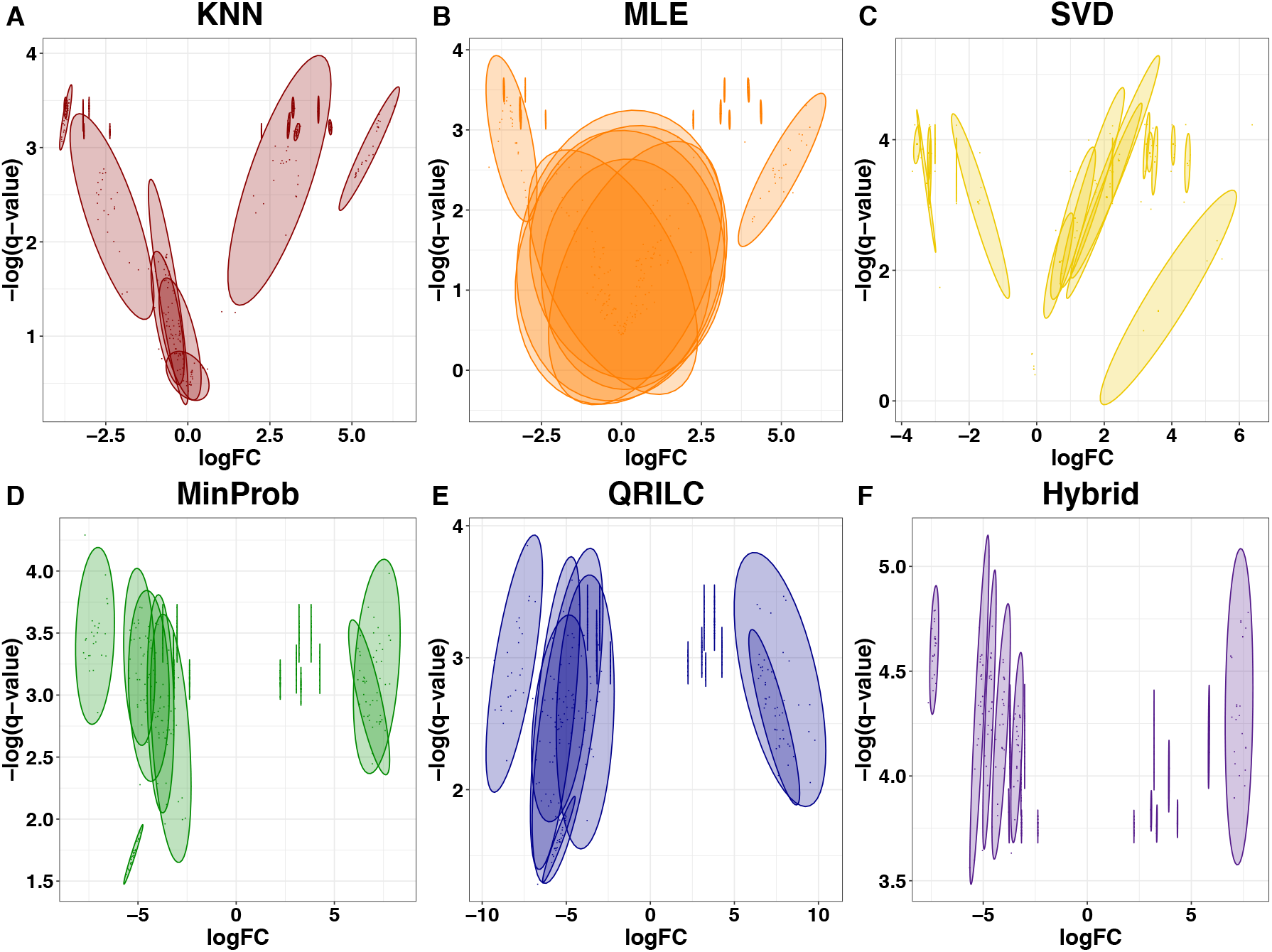
Spread plots of -log *q*-value vs logFC for merged top proteins across all imputation methods for MDA-MD-468 data. Following 25 consecutive iterations, each dataset from each imputation method was ranked by q-value mean. The top 10 proteins from each method was pooled into a top protein list (**Supplemental Table 2**) and plotted here. The ellipse represents a 95% confidence interval.

We merged the top 10 proteins from each method to a single list and constructed spread plots. This list contained 19 total proteins with the following distribution of missing values: 0 missing values – 11 proteins, 1 missing value - 1 protein, 2 missing values - 1 protein and 3 missing values - 6 proteins. The SFI-hybrid and MNAR methods performed better than the MAR **(Fig. 4 and Supp. Table 2).** The MLE method performed the worst, with *k*NN and SVD falling in the middle **(Supp. Table 2)**.

### Imputation influences with increasing number and type of missing values

The PXD004462 sample set consists of NTERA2 (NT2) pluripotent embryonic carcinoma cells treated with retinoic acid for 8 days to induce neuronal differentiation. Therefore, we expect the nuclei that were isolated for immunoprecipitation tandem mass spectrometry (IP-MS/MS) with EZH2 (n =3), SUZ12 (n = 3) and IgG (n = 3) to have a true presence/absence in protein profiles. The number of missing values in this dataset was more typical of bottom-up proteomics experiments, ranging from 6 – 50% (**Fig. 2B**). Data was ranked by *q*-value mean following multiple imputations and classified as described in the methods above.

Following removal of contaminants, reverse decoys and lowly expressed proteins, we identified 601 total proteins in the EZH2 IP. With a q-value threshold of 0.05, DEP lists included: 338 significant proteins when using *k*NN imputation, 348 for MLE, 375 for SVD, 326 for MinDet, 179 for MinProb, 198 for QRILC and 365 for SFI-hybrid. The SUZ12 IP contained 350 proteins after filtering and the resulting DEP lists included: 119 significant proteins when using *k*NN, 97 for MLE, 102 for SVD, 127 for MinDet, 120 for MinProb, 111 for QRILC and 187 for SFI-hybrid.

Approximately one-half of the proteins determined to be significant contained all observations (no missing values) in every imputation method for both IP experiments (**Supp. Figs. 5 and 9**). Increasing number of missing values resulted in a wider range of logFC protein expression values across imputation methods for the EZH2 and SUZ12 IPs (**Supp. Figs. 3 and 7**). We merged the top 10 proteins from each method to a single list and constructed spread plots. The merged top protein list for EZH2 IP consisted of 21 proteins (3 missing values – 3 proteins, 2 missing values – 3 proteins, 1 missing value – 3 proteins, and 0 missing values – 12 proteins) while the SUZ12 IP list contained 23 total proteins (3 missing values – 11 proteins, 1 missing value – 4 proteins, and 0 missing values – 8 proteins). The SFI-hybrid method performed better with increasing number of missing values as evidenced by the narrow ellipses in the spread plots showing the tighter ranges of values for LogFC (**Fig. 5 and Supp. Fig. 6)** and - log10 *q*-values (**Supp. Figs. 4 and 7**). These data illustrate the impact the missingness and appropriate choice of imputation method can have on the range of values obtained in DEP analysis. The combined protein ranks were determined as described in the methods section and used to evaluate the relative rank of each of the canonical PRC2 complex members from both the EZH2 and SUZ12 IPs. These rankings show that the SFI-hybrid method outperformed all other imputation methods, followed by the MNAR methods (MinDet, MinProb, QRILC), SVD, *k*NN and MLE (**Fig. 6A, 6B, 6D and Supp. Table 1**).

**Figure 5.**
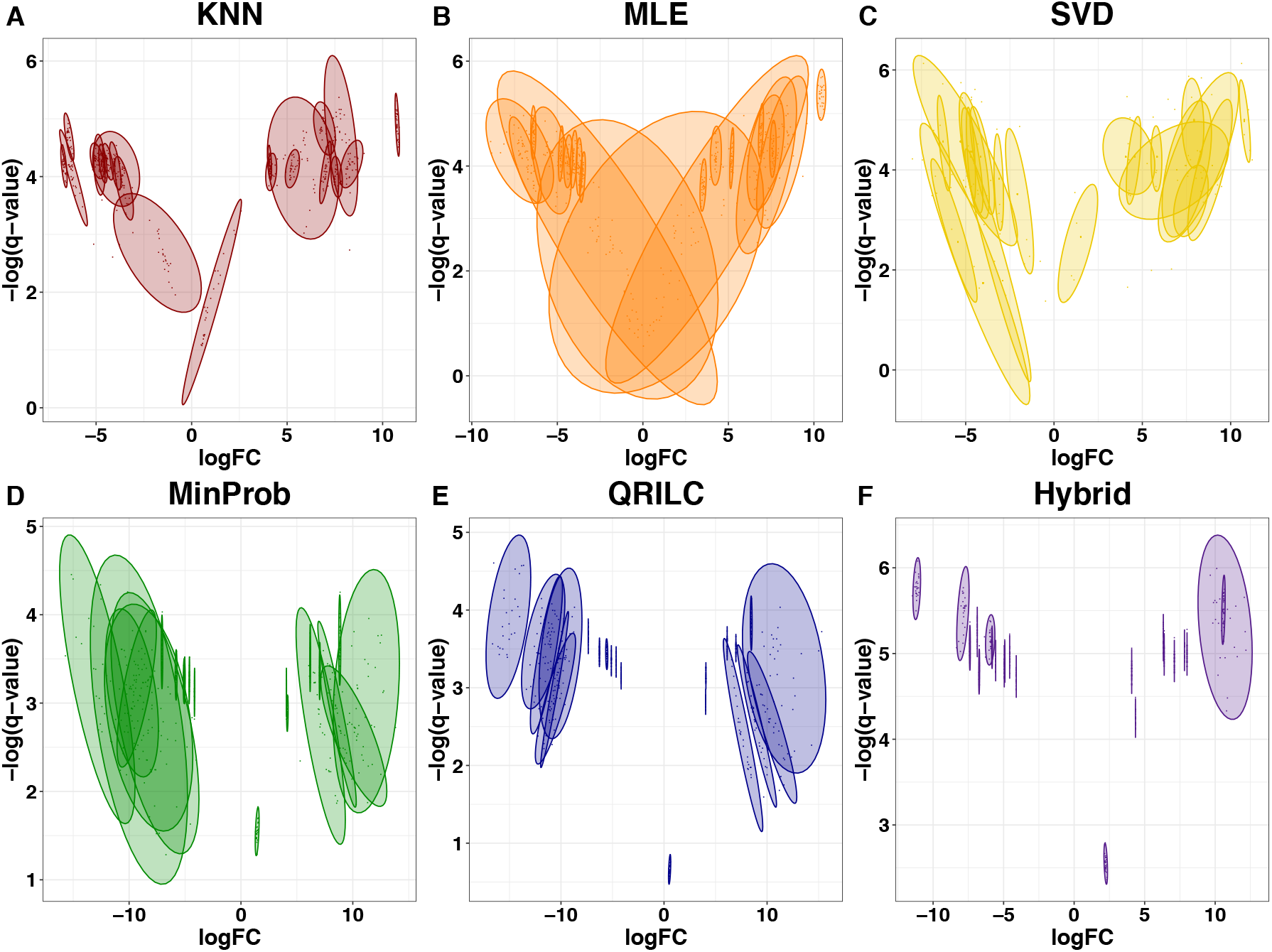
Spread plots of -log *q*-value vs logFC for merged top proteins (**Supplemental Table** 3) across all imputation methods for EZH2 IP data. Data was processed as described in Figure 4.

**Figure 6.**
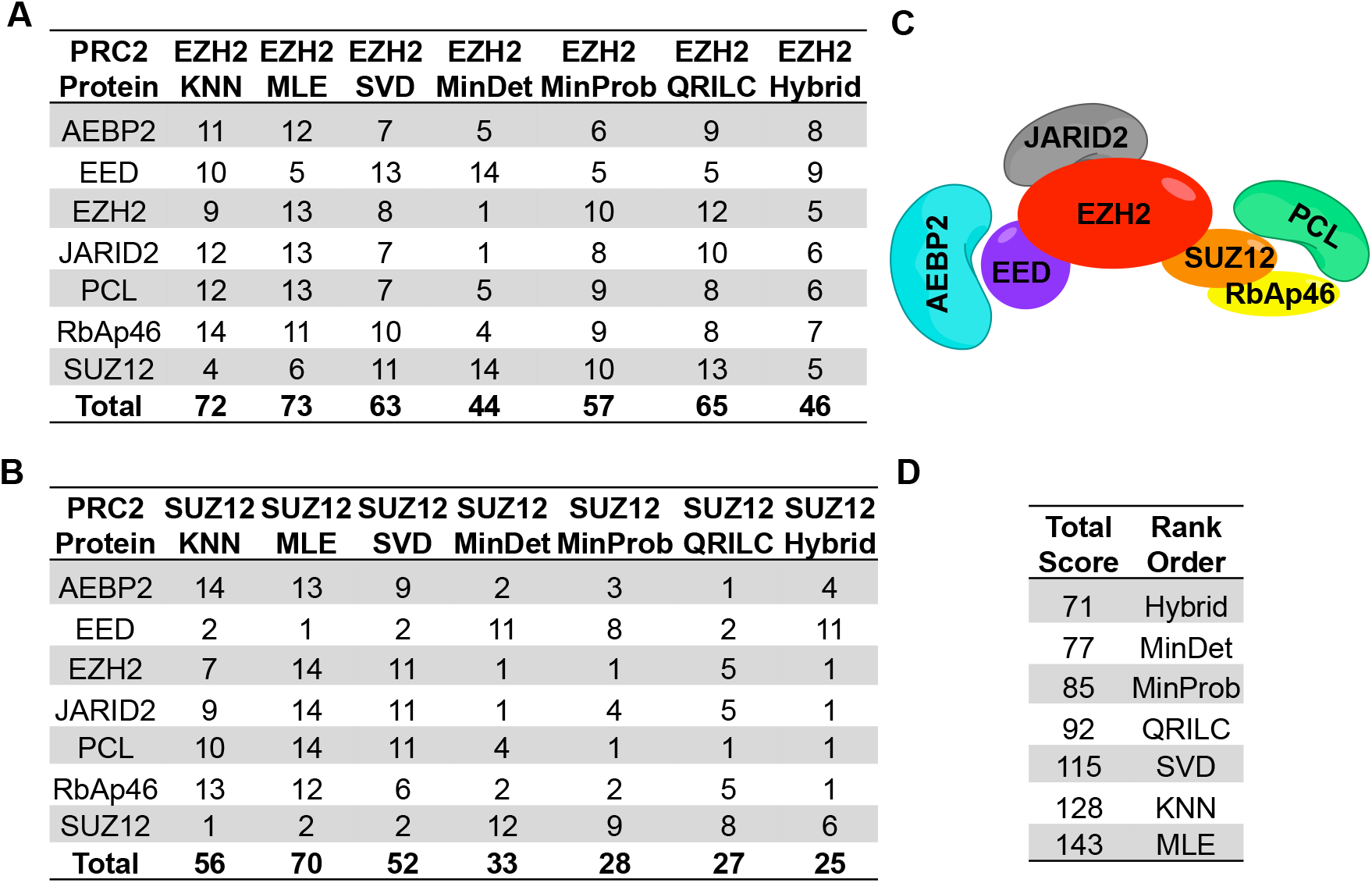
Relative rank ordering of PRC2 component IPs. Each component of the PRC2 complex identified in EZH2 (**B**) and SUZ12 (**C**) IPs was scored (1-14) according to position in the rank-ordered protein lists (**Supplemental Table 1**). (**C**) Canonical human PRC2 protein complex, adapted from Margueron, R., 2011, *The Polycomb Complex PRC2 and its Mark in Life.* Nature, 2011. **469**(7330): p. 343-9 (**D**) Total scores and overall ranking of each method.

## DISCUSSION

We investigated multiple imputation methods to combat the missing value problem in bottom-up proteomics. We chose to work with the methods within the imputeLCMD package because they span a wide range of imputation methods in an accessible R package (described in more detail in [11]). As described here, we have modeled two dissimilar sets of data to be all or mostly MAR, MNAR or a combination of the two. In the case of all MAR, the model.selector argument in the impute.MAR function was converted such that all rows in the dataset would be flagged strictly MAR. Our aim was to develop a robust technique that would be applicable across many types of data with transparency, accuracy and reproducibility.

Here, we propose utilizing a combinatorial MAR/MNAR, or SFI-hybrid, with a typical model selector but performing imputation separately for each treatment or group of data. This approach is shown to most accurately model data regardless of type. This observation is not unexpected in that even for highly similar data sets with replication and few missing values, the left-censored nature of proteomics data is well suited to hybrid imputation methods.

A meta-analysis of the MDA dataset revealed the SFI-hybrid method maintained the same level of significance across all missing value types with the smallest standard deviations **(Supp. Figs. 1 and 2)**. Additionally, there was a negative trend toward non-significance with increasing number of missing values in all methods excluding the SFI-hybrid **(Supp. Fig. 2)**. To examine the top significant proteins in the MDA dataset, we binned each protein into the number (0, 1, 2, 3) and type of missing value (B for missing in both treatments). As expected, fewer proteins were designated as significant when there were missing values present in both treatment groups **(Fig. 3)**. We found all methods, other than the SFI-hybrid, favored complete cases with greater than 60% of significant identifications containing no missing values. This observation suggests either the methods have a bias to choose complete cases or the algorithms are imputing values too close to the observed to be considered significant.

To further investigate the performance of the missing value imputation methods for preserving accurate logFC and *q*-values, spread plots were constructed as described in the results and methods sections. The SFI-hybrid and MNAR methods performed better at preserving the significance level than the MAR **(Fig. 4 and Supp. Table 2)** as all proteins were found to be significant and the confidence ellipses representing the standard deviation were located above the threshold cut-off (-log10 (*q*-value) > 1.3). This is expected as MNAR methods are designed specifically for the low-abundant nature of these absences and impute the left-censored data appropriately. The MLE method appears to have performed the worst, as six of the 19 proteins were not significant **(Supp. Table 2)**. Upon closer examination of the data, this only occurs when there are three missing values and can be explained by the large standard deviation after multiple imputations. The four proteins not significant with the *k*NN method and with opposite logFC values in the SVD method were also from the three missing value type **(Supp. Table 2)**. When investigating proteins with three missing values that are imputed using *k*NN or SVD, we observe that logFC is highly variable and can change direction as well. We would caution the choice to use a single imputation strategy with these two methods and, instead, encourage the use of 15-16 nearest neighbors with *k*NN and 6-7 principal components with SVD where the data calculations are far more stable. Altogether, these results imply the missing data is not all MAR or all MNAR and imputation should be performed with a strategy to reflect that.

Meta-analysis was performed with EZH2 and SUZ12 IPs as mentioned above with the MDA-MB-468 data). The SFI-hybrid method maintained similar level of significance across all missing value types with the smallest standard deviations in both IPs **(Supp. Figs. 4 and 8)**. The range of logFC values varied in the SUZ12 IP **(Supp. Fig. 7)** and is attributed to the large amount of missingness when compared to EZH2 IP. The more stringent SUZ12 IP resulted in fewer observed values leading to under-represented variation after imputation. Interestingly, there was an opposing trend with increasing missingness in the EZH2 IP; logFC values shrank towards zero when using MAR methods and expanded when imputing with MNAR methods **(Supp. Fig. 3)**. All imputation methods, excluding the SFI-hybrid, trended toward non-significance and large variance across *q*-value as missingness increased **(Supp. Figs. 4 and 8)**. As mentioned above, these observations offer further support that the algorithms for MAR methods are imputing values too close to the observed to be considered significant or have an inherent bias to choose complete cases.

Once the top 100 significant proteins in the IPs were binned according to missingness, we found the SFI-hybrid significant protein list evenly distributed across missingness with approximately 40% in complete cases or 3 missing values **(Supp. Figs. 5 and 9)**. In the EZH2 IP, all methods excluding the SFI-hybrid favored complete cases. Additionally, all other methods except MLE were able to determine significance when 3 missing values were present **(Supp. Fig. 5)**. In the case of SUZ12, the *k*NN and MLE MAR methods favored complete cases while SVD and MNAR modeled the same trend as the SFI-hybrid in the EZH2 IP **(Supp. Fig. 9)**.

All methods were able to call significance equally in the top protein list for EZH2 IP **(Supp. Table 3)**. However, the SFI-hybrid method is most consistent with imputing values for both sets of IPs as seen by the tight confidence ellipses **(Fig. 5 and Supp. Fig. 6)**. The *k*NN and MLE methods were the worst performers with the SUZ12 IP data; the top protein list generated as described in the methods section was characterized by the largest variances **(Supp. Fig. 6)** and the greatest number of non-significant protein identifications at 7 and 11 when imputing with these approaches **(Supp. Table 4)**. Additionally, 2 of the 7 and 5 of the 7 PRC2 complex proteins were determined as not significantly enriched in the *k*NN and MLE methods. It is interesting to note that these were instances where 3 missing values occurred. These observations suggest the missing data is not all MAR or all MNAR and caution should be taken to choose an imputation strategy that appropriately models the data such as the MI SFI-hybrid approach.

To determine the overall performance of the methods presented in this study, we focused on the IP dataset since it is more representative of a bottom-up proteomics experiment with values MNAR. We decided to examine the data from both IPs in a combinatorial manner because we did not want the results biased from the large range of missing values or the non-specificity of the EZH2 antibody **(Fig. 2B).** The canonical PRC2 complex **(Fig. 6C)** consists of seven proteins: AEBP2, EED, EZH2, JARID2 (JARD2), RbAp46 (RBBP7), SUZ12 and PCL family. We arbitrarily chose the PCL protein MTF2 for our ranking analysis. Following the MI strategy and ranking by *q*-value mean, we recorded the global rankings for each of the PRC2 components for each IP across each method separately **(Supp. Table 1)**. Relative ranking of the methods in each IP demonstrated MinDet performed best followed closely by SFI-hybrid for EZH2; the MNAR and SFI-hybrid methods outperformed the MAR methods when dealing with large amounts of missing data as in the SUZ12 IP **(Fig. 6A, 6B)**. Overall, we determined the SFI-hybrid had the best ranking (lowest score) of the canonical PRC2 components, followed by MNAR and MAR methods **(Fig. 6D)**.

In summary, we have explored MAR, MNAR and SFI-hybrid missing value imputation strategies used in intensity-based proteomics workflows. We have evaluated the performance of these methods while considering the extreme ends of missing values encountered in this type of bottom-up proteomics data analysis. In data with similar proteomic profiles, imputing missing values as all MAR or MNAR may be acceptable approaches when the nature of missingness is known or with complete case analysis (excluding situations of presence/absence). To avoid unnecessarily excluding data from an analysis, we propose a SFI-hybrid technique that demonstrates accuracy, reproducibility with resistance to the large number of missing values that influences the other methods investigated here.

## ABBREVIATIONS

DEP: differentially expressed proteins
DDA: data-dependent acquisition
FDR: false discovery rate
IP: immunoprecipitation
IP-MS/MS: immunoprecipitation tandem mass spectrometry
*k*NN: *k-*nearest neighbors
LFQ: label-free quantitation
LOD: limit of detection
MAR: missing at random
MCAR: missing completely at random
MI: multiple imputation
MinDet: deterministic minimum
MI-MFA: multiple imputation in multi-factor analysis
MinProb: probabilistic minimum
MLE: maximum likelihood estimation
MNAR: missing not at random
PRIDE: PRoteomics IDEntifications
QRILC: quantile regression imputation of left-censored data
SVD: singular value decomposition

## DISCLOSURE

Michael A. Freitas is the co-owner, co-founder, and Chief Scientific Officer of MassMatrix Inc. a for profit biotech company focusing on development of bioinformatics software for biopharma. The work in the publication are the sole work product of the authors and the Ohio State University and not a product of MassMatrix Inc.

## FUNDING ACKNOWLEDGEMENTS

Research reported in this manuscript was supported by the Comprehensive Cancer Center of The Ohio State University and by grants from the U.S. National Institute of Health/National Cancer Institute and National Institute of General Medical Sciences (Award Numbers P30CA016058 and R01GM122436).

