## Supplemental_Figures for "Multiple Imputation Approaches Applied to the Missing Value Problem in Bottom-up Proteomics"

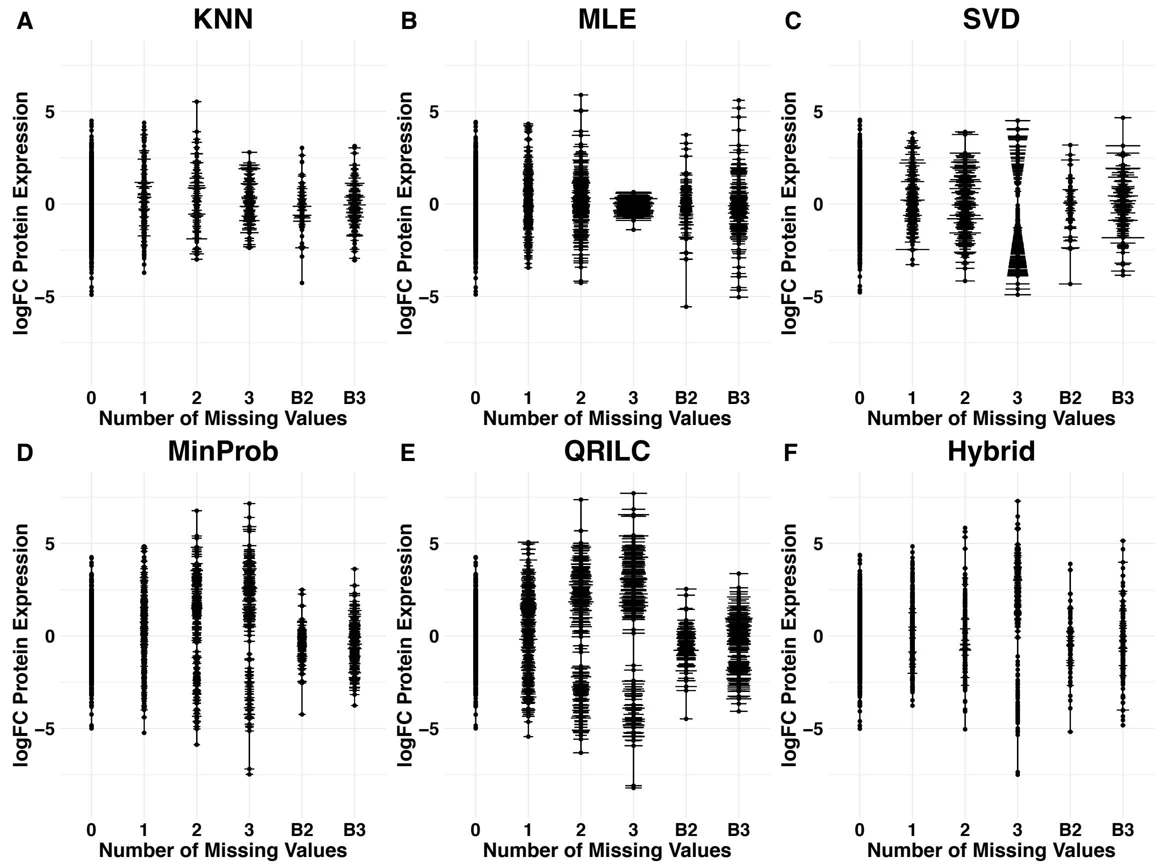


**Supplemental Figure 1.** Range of logFC protein expression as a function of the number and type of missing values in MDA-MB-468 following 25 consecutive iterations of each imputation method. The number of missing values can be missing in one sample group as 0, 1, 2 or 3 or in any combination of both sample groups as B2 or B3. The horizontal lines represent the standard deviation in the logFC values across the multiple imputations scaled by a factor of 0.5.

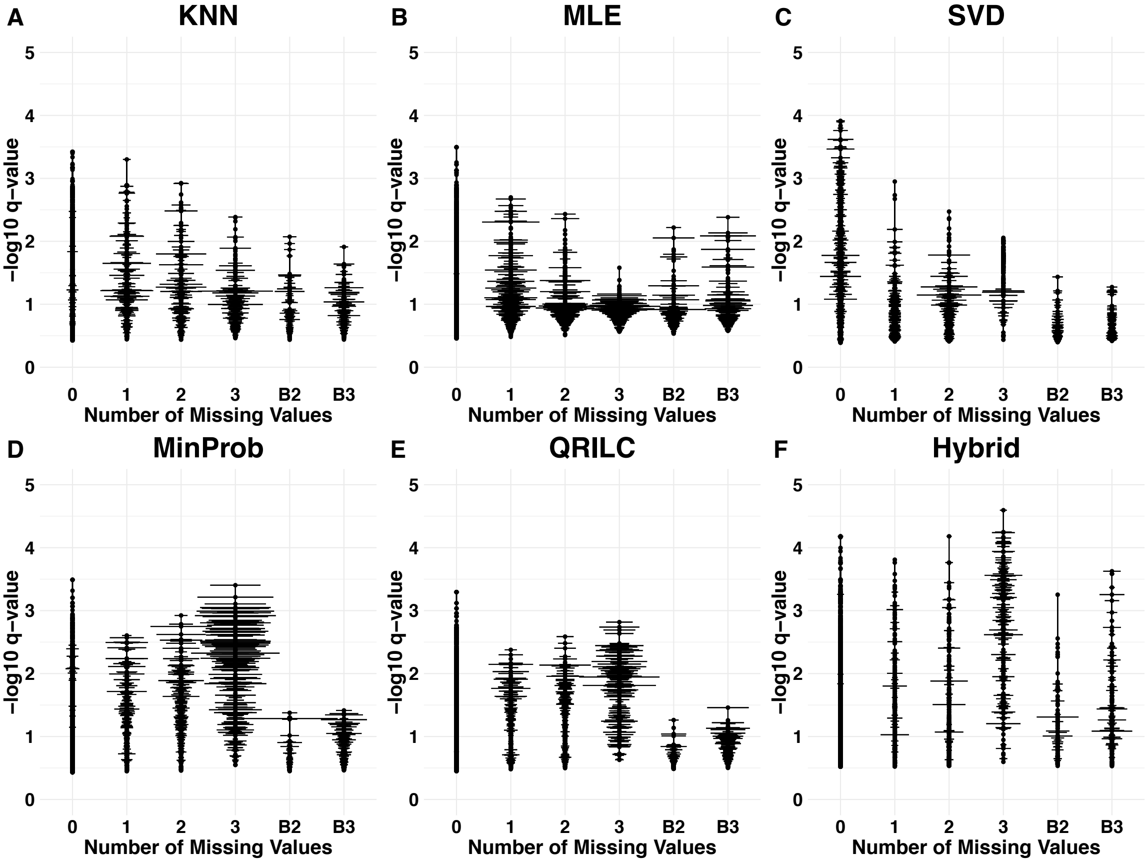


**Supplemental Figure 2.** Range of -log10 *q*-value mean as a function of the number and type of missing values in MDA-MB-468 following 25 consecutive iterations of each imputation method. The number of missing values can be missing in one sample group as 0, 1, 2 or 3 or in any combination of both sample groups as B2 or B3. The horizontal lines represent the range of *q*-value means across the multiple imputations scaled by a factor of 0.5.

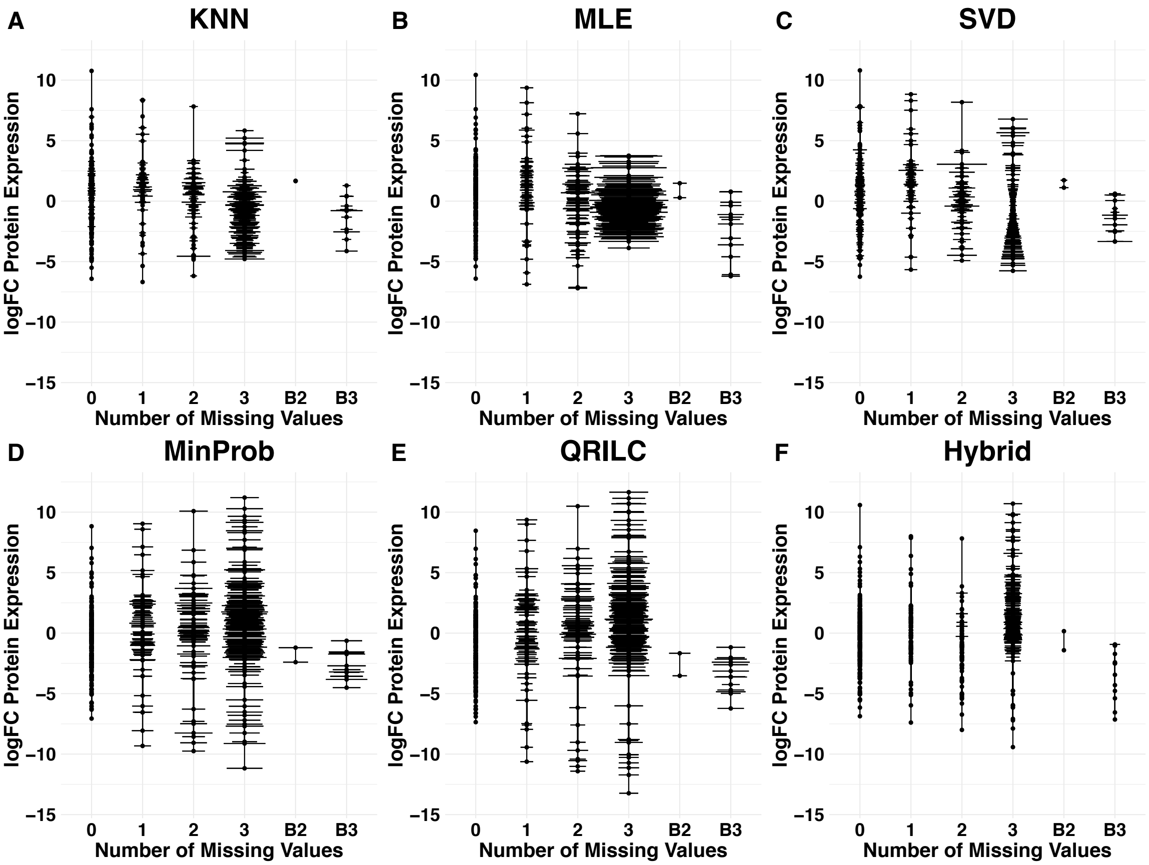


**Supplemental Figure 3.** Range of logFC protein expression as a function of the number and type of missing values in EZH2 IP compared to IgG control. Data was processed as described in Supplemental Figure 2.

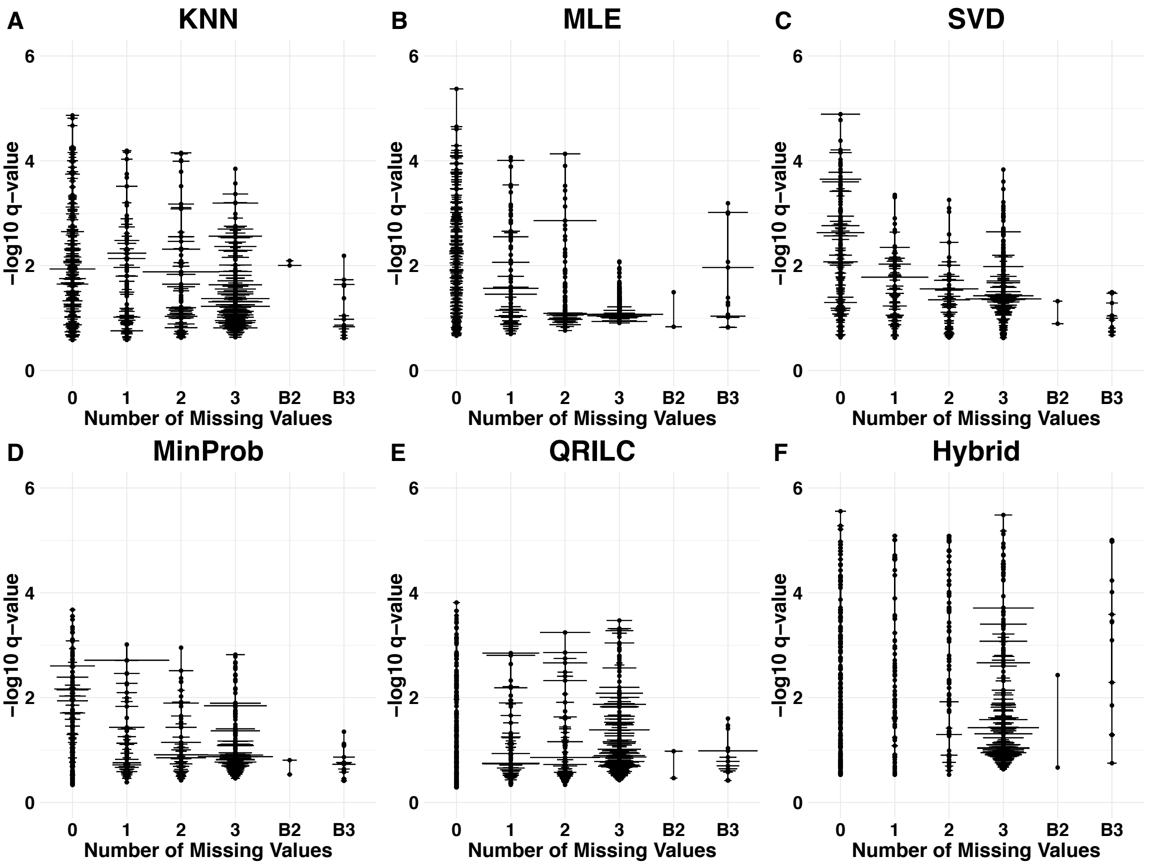


**Supplemental Figure 4.** Range of -log10 *q*-value mean as a function of the number and type of missing values in EZH2 IP compared to IgG control. Data was processed as described in Supplemental Figure 3.

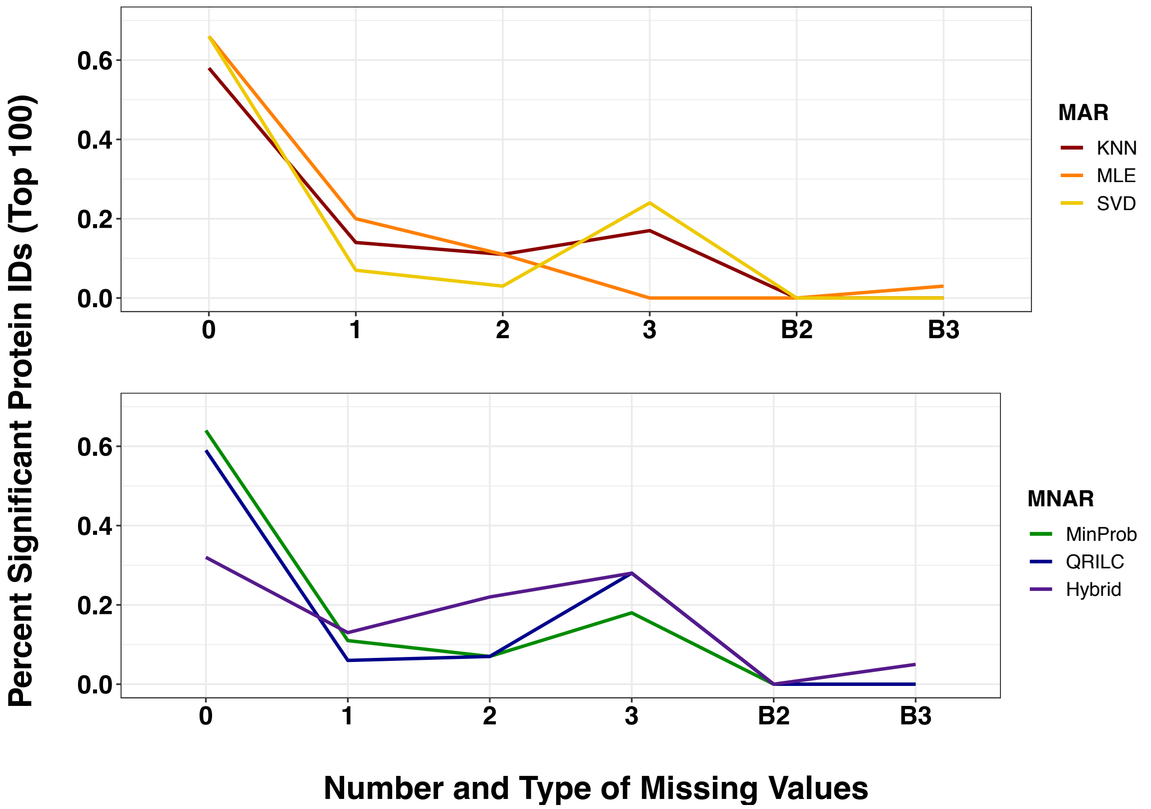


**Supplemental Figure 5.** Distribution of missingness across all imputation methods with the top 100 significant proteins identified in EZH2 IPs (n = 3). Data was processed as described in Figure 3.

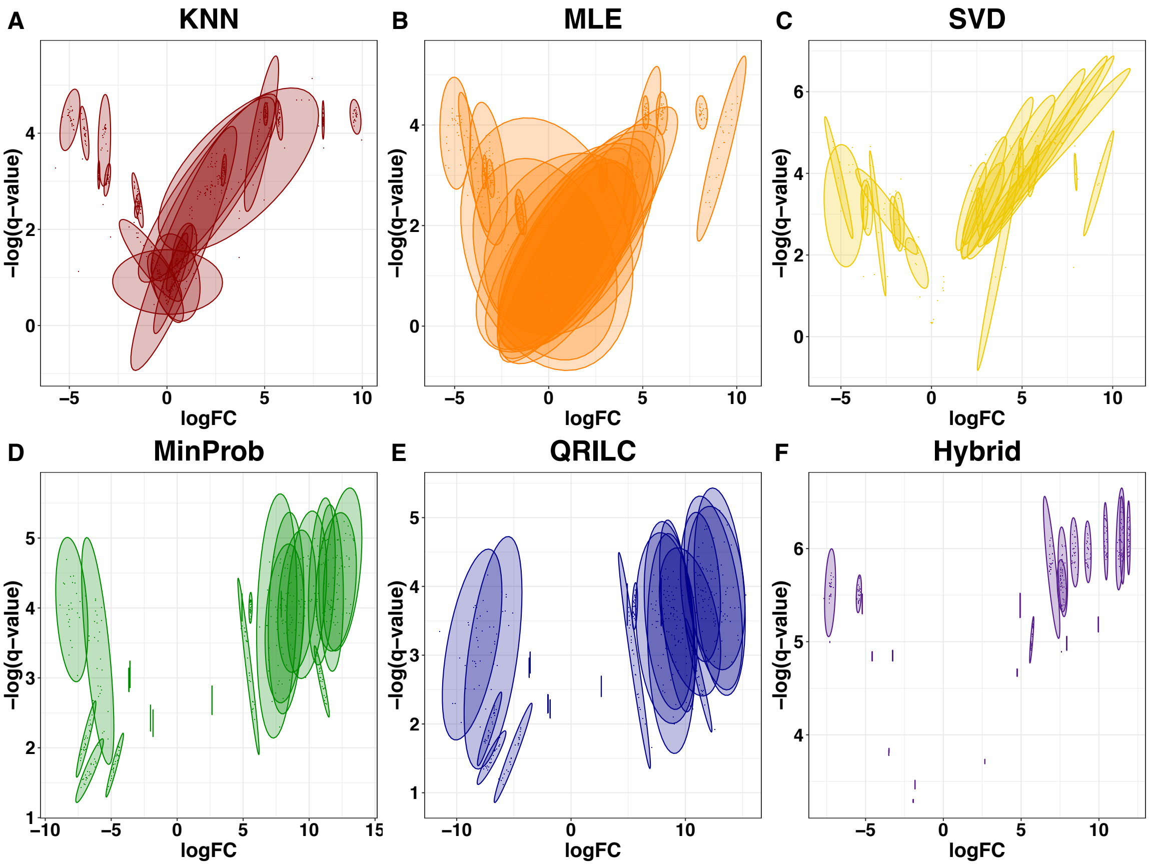


**Supplemental Figure 6.** Spread plots of -log *q*-value vs logFC for merged top proteins (**Supplemental Table 4**) across all imputation methods for SUZ12 IP data. Data was processed as described in Figure 4.

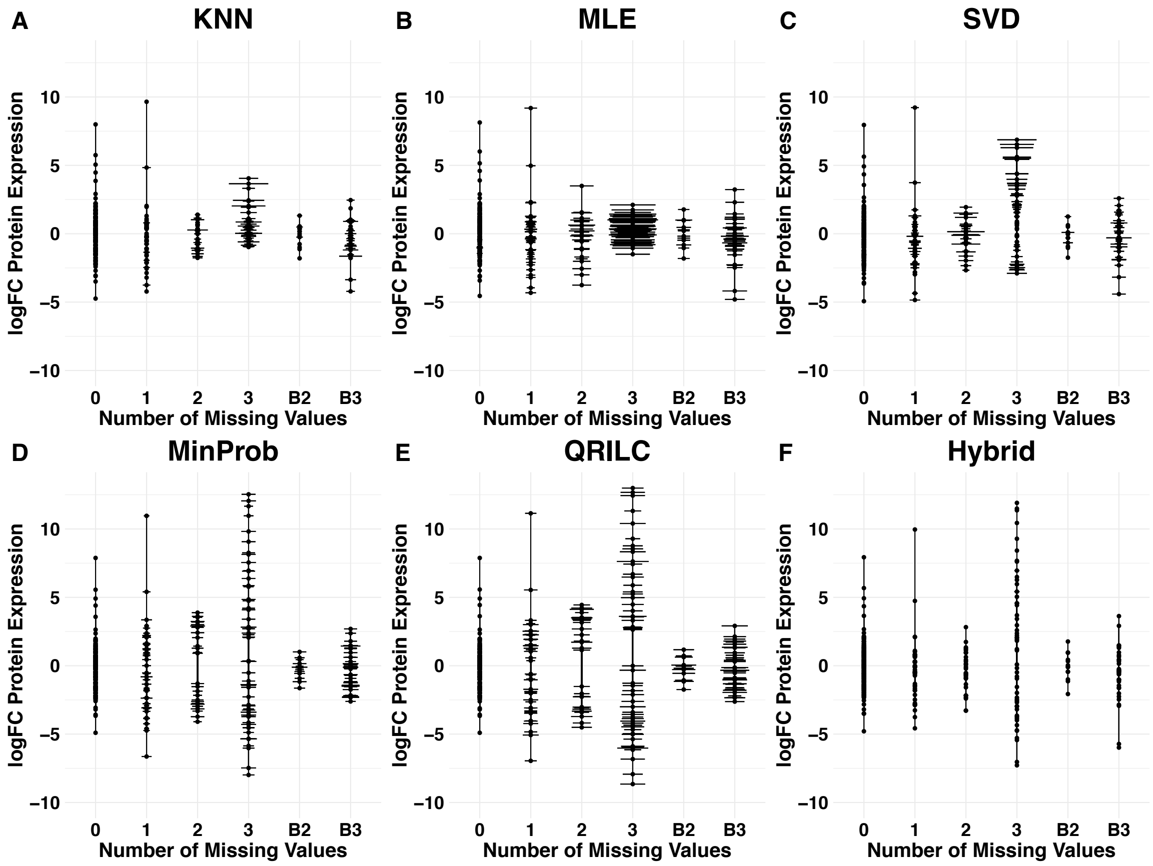


**Supplemental Figure 7.** Range of logFC protein expression as a function of the number and type of missing values in SUZ12 IP compared to IgG control. Data was processed as described in Supplemental Figure 2.

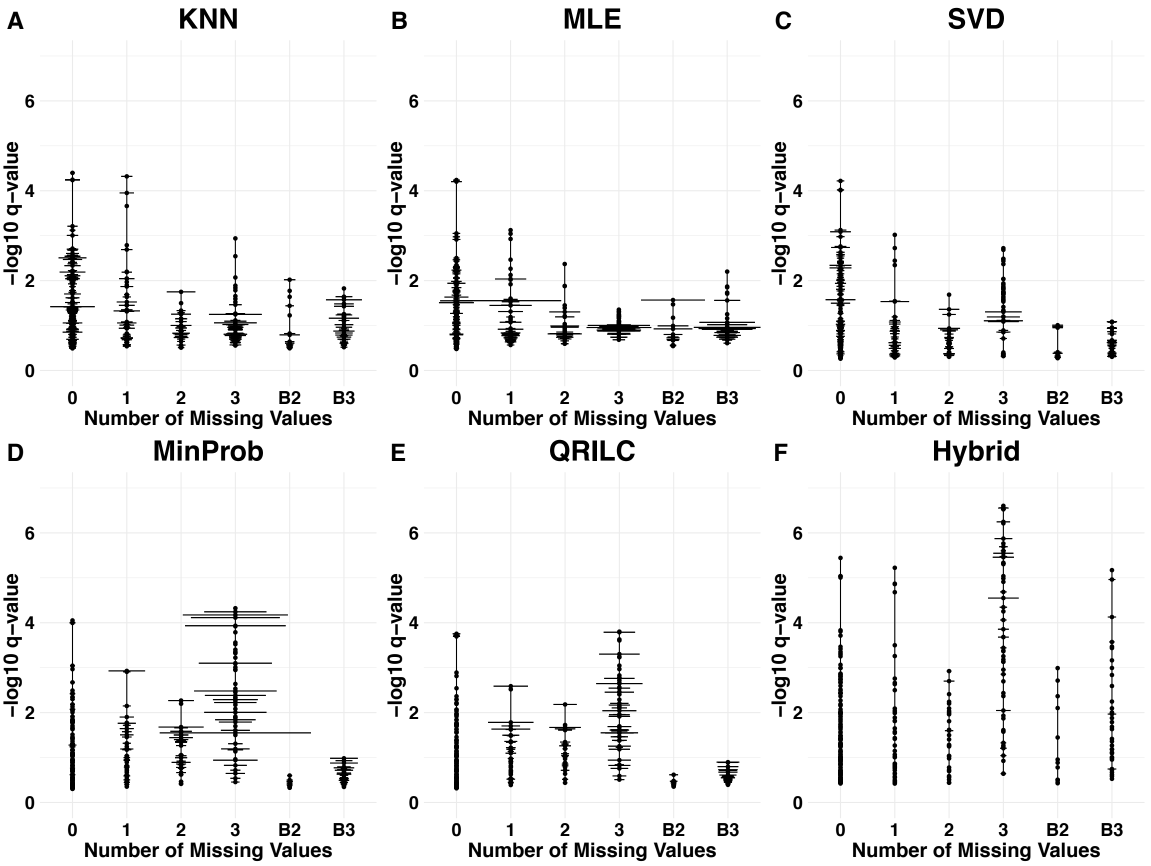


**Supplemental Figure 8.** Range of -log10 *q*-value mean as a function of the number and type of missing values in SUZ12 IP compared to IgG control. Data was processed as described in Supplemental Figure 3.

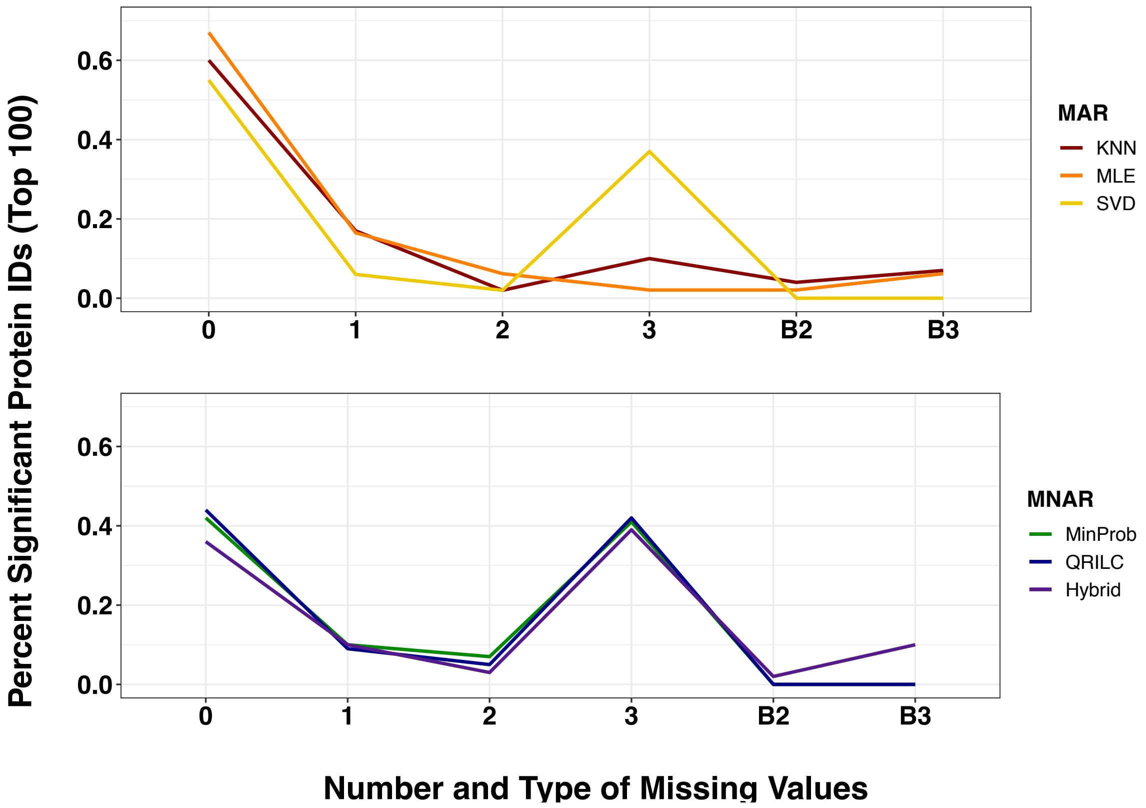


**Supplemental Figure 9.** Distribution of missingness across all imputation methods with the top 100 significant proteins identified in SUZ12 IPs (n = 3). Data was processed as described in Figure 3.

**A**

| PRC2 Protein | EZH2 KNN | EZH2 MLE | EZH2 SVD | EZH2 MinDet | EZH2 MinProb | EZH2 QRILC | EZH2 Hybrid |
|---|---|---|---|---|---|---|---|
| AEBP2 | 62 | 158 | 42 | 6 | 35 | 56 | 46 |
| EED | 20 | 4 | 33 | 38 | 4 | 4 | 18 |
| EZH2 | 24 | 176 | 12 | 1 | 26 | 30 | 6 |
| JARID2 | 125 | 216 | 18 | 3 | 34 | 44 | 17 |
| PCL | 127 | 157 | 24 | 7 | 42 | 29 | 21 |
| RbAp46 | 455* | 328 | 150 | 13 | 76 | 42 | 41 |
| SUZ12 | 8 | 18 | 34 | 120 | 31 | 78 | 10 |

**B**

| PRC2 Protein | SUZ12 KNN | SUZ12 MLE | SUZ12 SVD | SUZ12 MinDet | SUZ12 MinProb | SUZ12 QRILC | SUZ12 Hybrid |
|---|---|---|---|---|---|---|---|
| AEBP2 | 163* | 128* | 56 | 2 | 3 | 1 | 4 |
| EED | 3 | 1 | 3 | 22 | 5 | 3 | 22 |
| EZH2 | 10 | 125* | 27 | 1 | 1 | 6 | 1 |
| JARID2 | 43 | 162* | 61 | 3 | 4 | 7 | 3 |
| PCL | 58 | 135* | 63 | 4 | 2 | 2 | 2 |
| RbAp46 | 209* | 173* | 20 | 12 | 12 | 16 | 9 |
| SUZ12 | 2 | 6 | 6 | 41 | 26 | 22 | 18 |

**Supplemental Table 1.** Global rankings of proteins identified in PRC2 IPs. EZH2 (**A**) and SUZ12 (**B**) IPs were rank-ordered by *q*-value mean for each imputation method following consecutive iterations and the final position out of all protein identifications was recorded.

* designates non-significance at a *q*-value threshold cutoff < 0.05.
